# The computational models of AlphaFold2 and RoseTTAfold carry protein foldability information

**DOI:** 10.1101/2022.01.27.477978

**Authors:** Sen Liu, Kan Wu, Cheng Chen

## Abstract

Protein folding has been a “holy grail” problem of biology for fifty years, and the recent breakthrough from AlphaFold2 and RoseTTAfold set a profound milestone for solving this problem. AlphaFold2 and RoseTTAfold were successful in predicting protein structures from peptide sequences with high accuracy. Meanwhile, although the protein folding problem also cares about the kinetic pathways of protein folding, AlphaFold2 and RoseTTAfold were not trained with this functionality. Considering that their training sets contain proteins from foldable sequences and sequence evolutionary information, we wondered if the computational models from AlphaFold2 and RoseTTAfold might carry protein foldability information. To test this idea, we systematically predicted the structural models of 149 circular permutants and 148 alanine insertion mutants of the 149-residue dihydrofolate reductase of Escherichia coli with AlphaFold2 and RoseTTAfold. Our data showed that although AlphaFold2 and RoseTTAfold could not directly identify unfoldable proteins, the structural variations of computational models are correlated with protein foldability. Furthermore, this correlation is independent of secondary structures. Most importantly, the structural variations of computational models are quantitatively correlated with protein foldability but not protein function. Our work could be of great value to the design of circular permutants, the design of fragment complementary proteins, the design of novel proteins, and the development of computational tools for predicting protein folding kinetics.

**Highlights:**

1. AlphaFold2 and RoseTTAfold could not directly identify unfoldable proteins.
2. The structural variations of computational models are correlated with protein foldability.
3. This correlation is independent of secondary structures.
4. The structural variations of computational models are quantitatively correlated with protein foldability but not protein function.

## Introduction

Proteins are synthesized as linear chains of amino acids, but they generally could not perform biological functions before forming specific three-dimensional (3D) conformations. The formation of the 3D structure of a peptide sequence, i.e., protein folding, is tremendously challenging because the large dimensionality and its stochasticity. However, the seminal work by Christian Anfinsen and colleagues led to the hypothesis that the peptide sequence of a protein intrinsically determines its specific 3D structure ^1^. Since then, protein folding has been described as a searching for the lowest-energy conformation in the energy landscape ^2^. This hypothesis further led to the extensive investigation of the protein folding problem, which includes three questions: what is the protein folding code, how to predict the 3D structure of a given peptide sequence, and what are the kinetic folding pathways of proteins ^2,3^.

With the accumulated contributions of many scientists, remarkable achievements have been made in solving the protein folding problem in the last fifty years ^2^. The biggest breakthrough was recently achieved by AlphaFold2 ^4^ and RoseTTAfold ^5^. Both methods took advantages of previous knowledges on protein folding and the recent development of computational algorithms and hardware. By incorporating physical constraints, evolutionary information, neural networks, and GPU computing, AlphaFold2 and RoseTTAfold were able to predict protein structures comparable to experimental accuracy. Therefore, these methods are extremely helpful when we want to know the possible 3D structure of a natural protein. However, a virtually predicted model is far from a viable folded protein. It is unknown yet whether a peptide chain could be really foldable when it is virtually folded. For example, David Baker and colleagues recently designed 129 proteins computationally with trRosetta but discovered that only 27 proteins folded well after expression ^6^.

Circular permutation is a protein engineering strategy to elucidate the structure-function relationship and folding kinetics of proteins ^7^. In circular permutation, the peptide chain of a protein is rearranged by joining the N-terminus and the C-terminus while new termini are generated by the cleavage of a peptide bond other than the original ones (Figure 1a). Therefore, circular permutation changes the connectivity but not the composition of residues of a protein. Besides being a research tool, circular permutation also occurs in natural proteins as an evolution strategy ^8^. Studies showed that certain circular permutation affects the folding progress of a protein while keeping the overall structure and function without significant changes ^7,9–11^. If the cleavage in a contiguous peptide segment results in the complete loss of the ability of the protein to fold, this sequence region is named as a folding element, which might play key roles in the formation of folding nuclei during the protein folding process ^11^. Previous studies proposed that the presence but not the order of folding elements is essential for a protein to be foldable ^9,11–13^. Although neither AlphaFold2 nor RoseTTAfold was designed to predict folding elements and folding kinetics, both methods took advantages of sequence evolution information. Therefore, it is of great interest to know if these state-of-the-art computational methods could distinguish foldable circular permutants from unfoldable ones of a natural protein.

**Figure 1.**
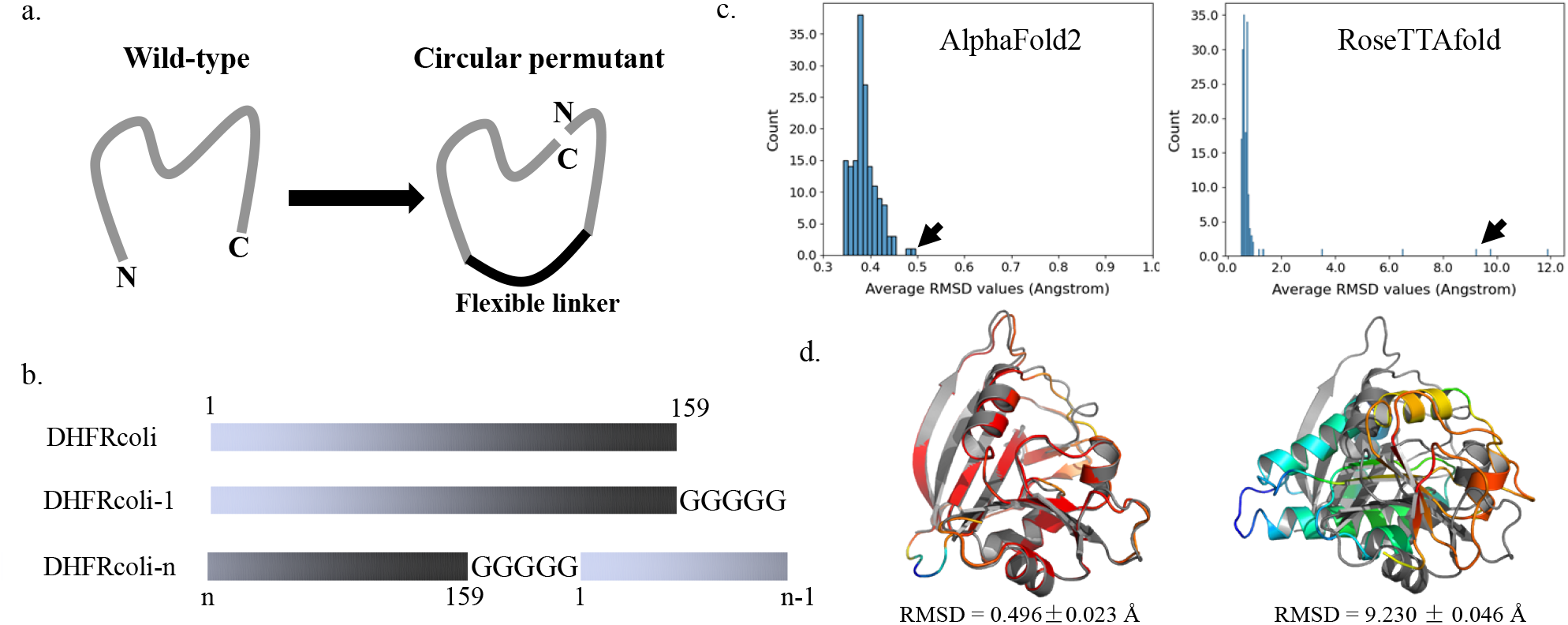
Construction of the circular permutants and model quality. (a) Scheme of circulate permutation. The original termini are connected by a flexible peptide linker, and new termini are introduced by the cleavage of a peptide bond elsewhere. (b) Scheme of the sequence construction of the DHFRcoli circular permutants. The marked numbers are residue numbers. A five-glycine linker was added to connect the original termini. (c) The distributions of the backbone RMSD values of the 159 AlphaFold circular permutants. The pointed models by the arrows are shown in D. (d) A predicted model from AlphaFold2 for the experimentally unfoldable circular permutant DHFRcoli-92, and A predicted model from RoseTTAfold for the experimentally foldable and active circular permutant DHFRcoli-79. The RMSD values were averaged from five models and calculated as the backbone RMSD aligned to the X-ray structure (PDB ID: 1RX4). The X-ray structure of the wild-type DHFRcoli is colored in gray and the models are colored as spectrum by pLDDT values of Cα atoms (red: higher pLDDT values; blue: lower pLDDT values).

The dihydrofolate reductase of Escherichia coli (EC 1.5.1.3; referred as DHFRcoli in this work) is an intensively studied model in protein circular permutation. By generating circular permutants at all 158 neighbored residue pairs of this 159-residue protein, Masahiro Iwakura and colleagues defined ten folding elements of DHFRcoli based on experimental data ^11^. Breaking any one of these folding elements abolished the correct folding of DHFRcoli. Interestingly, these folding elements did not strictly align with the secondary structural units, or the substrate and coenzyme binding sites ^11^. Nonetheless, the residues in these folding elements were correlated with the formation of the folding nuclei in the early folding events ^11^.

In this work, we systematically modelled the 3D structures of the 158 circular permutants of DHFRcoli with both AlphaFold2 and RoseTTAfold. Although not all permutants were experimentally foldable and active, they were all virtually folded and highly resembled the X-ray structure of the wild-type protein, indicating that these computational methods cannot evaluate the foldability of proteins. However, the conformational variations of the theoretical models of unfoldable permutants were larger than that of foldable permutants. Furthermore, the correlation between conformational variations and foldability was independent of secondary structures. Similar findings were discovered by modeling another 158 alanine insertion mutants of DHFRcoli ^14^ using both AlphaFold2 and RoseTTAfold. Our data indicated that the conformational variation of the computational model from AlphaFold2 and RoseTTAfold could be useful for evaluating protein foldability.

## Results

### Foldable and unfoldable circular permutants have folded 3D models from AlphaFold2 and RoseTTAfold

DHFRcoli is a 159-residue protein with well characterized crystal structures. As a control, we modelled the structure of the wild-type protein using AlphaFold2 and RoseTTAfold. The model quality was evaluated by the backbone RMSD value between the model and the X-ray structure (PDB ID 1RX4). The RMSD values of the AlphaFold2 model and the RoseTTAfold model were 0.369±0.022 Å and 0.529±0.003 Å respectively (Figure S1), indicating that both methods recaptured the native structure with high accuracy.

In the 158 circular permutants, a five-glycine linker was inserted between the original N-terminal and C-terminal residues ^11^. Therefore, this five-glycine linker was also appended to the C-terminus of the wild-type sequence to serve as the reference sequence. This linker was optimized and did not perturb the core structure and function of DHFRcoli according to the previous investigation ^10^ and an X-ray structure containing four glycine residues at the C-terminus (PDB ID: 5UII). Since this sequence also could be considered as an additional circular permutant with the first residue being the original N-terminus, it was referred as DHFRcoli-1. Accordingly, the circular permutant with the n^th^ residue as the N-terminus was named as DHFRcoli-n in this work (Figure 1b). Among the 159 permutants (including DHFRcoli-1), 86 were assigned as foldable and 73 were unfoldable based on the experimental data from trimethoprim (TMP) sensitivity assay, activity assay, and circular dichroism (CD) measurement ^11^ (Table S1). However, the RMSD values showed that AlphaFold2 gave all permutants 3D models closely resembling the wild-type structure (Figure 1c & 1d). Several models from RoseTTAfold had large RMSD values and were very different from the wild-type structure (Figure 1c), but some of them were actually foldable and active in experimental assays (Figure 1d & S2). This data indicated that, although they could generate computationally folded models for all DHFRcoli permutants, AlphaFold2 and RoseTTAfold were unable to classify foldable and unfoldable permutants. Therefore, whether AlphaFold2 or RoseTTAfold could generate well-folded models or not is not a reliable indicator for evaluating the actual foldability of a peptide sequence.

### Conformational variation correlates with the foldability of the circular permutants

Structural flexibility is a major cause of frustration in protein folding, and permutation could introduce or eliminate this kind of frustration, leading to the change of folding kinetics ^7^. A protein would be identified as unfoldable when it is kinetically trapped by frustration. We propose that if such frustration is introduced in to a DHFRcoli permutant, it might be possible to notice relatively large structural variations in these models. This hypothesis could be true for AlphaFold2 and RoseTTAfold models, because frustrated residue-residue contact information might be less represented in their training data based on experimental structures from foldable proteins and then less accurately predicted. Based on the experimental data, we noticed that the model quality from RoseTTAfold was much lower than AlphaFold2 and did not agree well with the experimental evidence (Figure 1c, 1d & S2), so we only used the AlphaFold2 models for further analyses. Indeed, we noticed that unfoldable permutants had larger structural variations (Figure 2a). Since DHFRcoli-1 only had five glycine residues appended to the C-terminus and minimally affecting the structure, we chose to use the RMSD value of DHFRcoli-1 as a threshold to split all permutants into two categories: the “High RMSD” group contains the permutants with RMSD values higher than that of DHFR-1, and the “Low RMSD” group contains the rest. Then it is noticeable that the Low RMSD group had more foldable permutants (59.8%) than the High RMSD group (43.9%) (Figure 2b). Meanwhile, the latter contained more unfoldable permutants than the former (56.1% v.s. 40.2%). A note is that the original paper ^11^ mentioned that some soluble (foldable) permutants were refolded in vitro from inclusion bodies, but the paper did not provide enough information that could be used to tell if and how many permutants in the High RMSD group were among them. In addition, the permutants with large RMSD values formed local clusters (Figure 2c), indicating that the structural variation was a result of circular permutation instead of random quality fluctuations of structural prediction.

**Figure 2.**
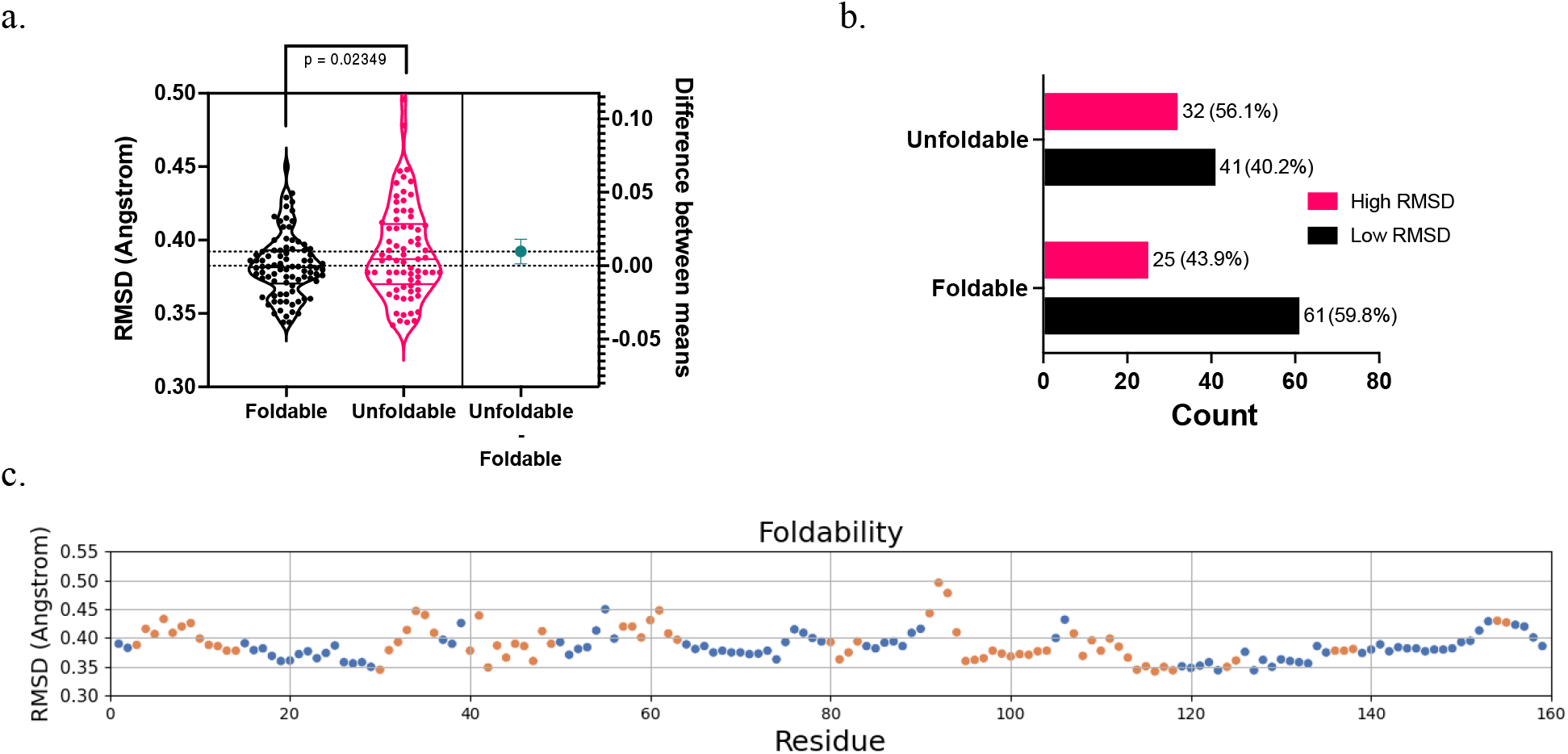
The RMSD variations of the AlphaFold2 models of DHFRcoli circular permutants. (a) The comparison of the model RMSD values of experimentally foldable and unfoldable permutants. The statistical p value was calculated with the unpaired t-test. (b) The amounts of the foldable and unfoldable permutants in the Low RMSD group and the High RMSD group. The numbers beside the bar are the permutant number and the percentage in the corresponding group. (c) The RMSD distribution of the 159 DHFRcoli circular permutants. The blue dots indicate the experimentally foldable permutants, and the orange dots indicate the unfoldable ones.

### The correlation between conformational variation and foldability is independent of structural elements

Both experimental and theoretical data support the formation of folding nuclei during protein folding ^15^. Based on the cleavage sites of unfoldable circular permutants of DHFRcoli, Iwakura et al. assigned the 73 corresponding DHFRcoli residues to ten “Folding Elements” that might play important roles in the formation of folding nuclei ^11^. Accordingly, the other 86 residues do not belong to Folding Elements (Figure 3a). Since secondary structure units (alpha helices and beta sheets) are the main structural elements of folded proteins, we asked if the High RMSD group contains more residues within secondary structure units than the Low RMSD group.

**Figure 3.**
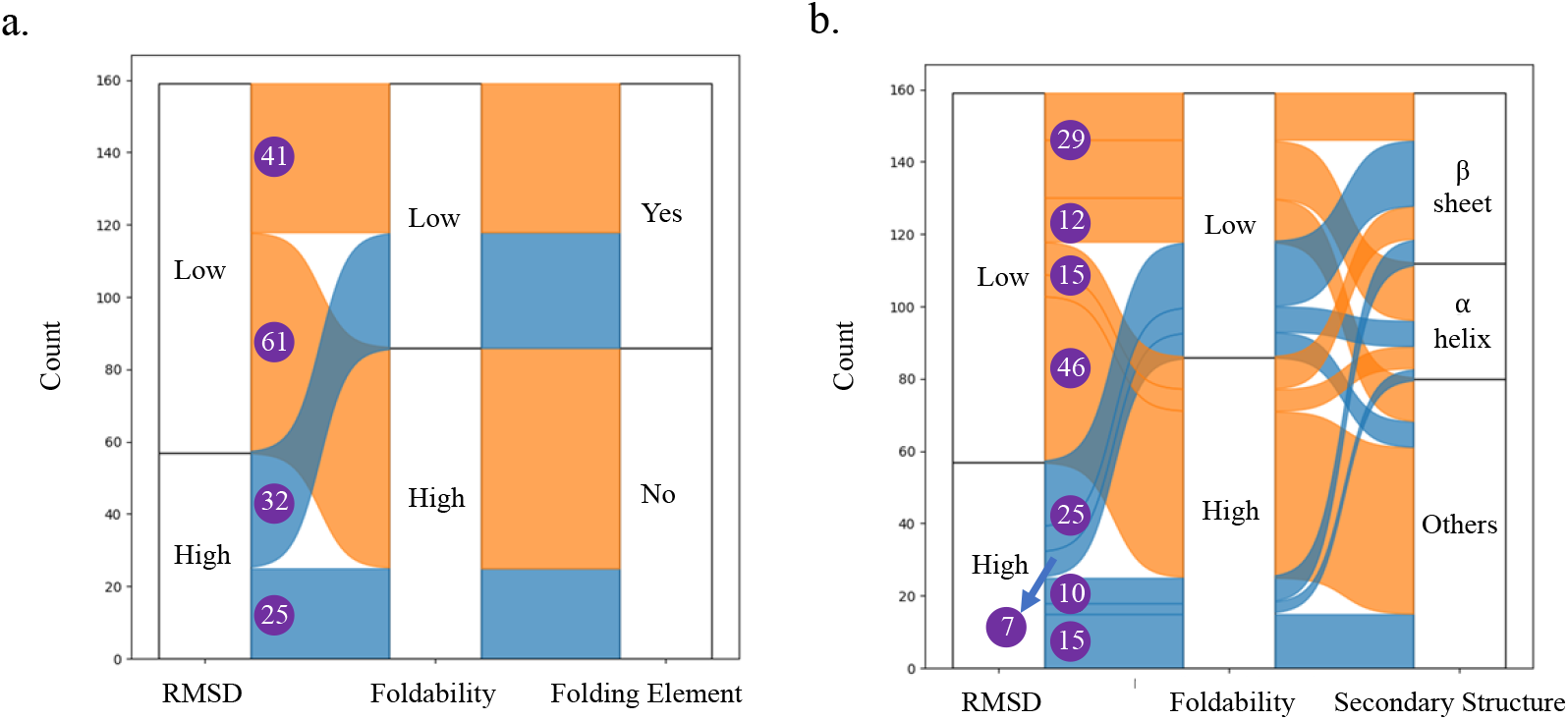
The correlation between the structural variations of the AlphaFold2 models of the 159 DHFRcolil circular permutants, their experimental foldability, and structural elements. (a) Iwakura et al. assigned the DHFRcoli residues into “Folding Elements” if the corresponding circular permutant had low foldability ^11^. (b) The correlation between RMSD values of the models, the foldability of the permutants, and the secondary structure units of the residues. The numbers in purple circles are the numbers of the permutants in different categories.

Among the 159 residues of DHFRcoli, 79 (49.7%) residues form secondary structure units (alpha helices and beta sheets), and 80 (50.3%) residues belong to random coils. As shown in Figure 3b, the 79 permutants with the cutting site in secondary structure units, 25 (31.6%) were foldable and 54 (68.4%) were unfoldable. The 80 permutants with the cutting site out of secondary structure units, 61 (76.2%) were foldable and 19 (23.8%) were unfoldable. Therefore, bond cleavage within secondary structure units resulted in unfoldable permutants with higher probability. When the cutting site was within secondary structure units, there were more foldable permutants in the Low RMSD group than that in the High RMSD group (15 v.s. 10). Similarly, when the cutting site was out of secondary structure units, there were also more foldable permutants in the Low RMSD group than that in the High RMSD group (46 v.s. 15). When the secondary structure rule and the model RMSD rule were combined, the percentage of foldable permutants in the Low RMSD group (34.1%, or 15/44) was higher than the percentage of the foldable permutants in the High RMSD group (28.6%, or 10/35). When the cutting site was out of secondary structure units, these percentages of foldable permutants were 79.3% (46/58) and 68.2% (15/22) respectively. Therefore, no matter where the cutting site is, the permutants in the Low RMSD group has higher foldability than those in the High RMSD group.

### Alanine insertion confirms the conclusions from circular permutation

To further investigate the folding of DHFRcoli, Shiba et al. ^14^ systematically constructed 158 alanine insertion mutants of DHFRcoli. They obtained the precipitant ratios of all mutants by comparing the fluorescence intensities of the protein bands on denatured SDS-PAGE gels. A mutant with a precipitant ratio (protein in supernatant/total protein) less than 60% was defined as foldable by Shiba et al. ^14^. In their design, an alanine residue was inserted between one pair of neighbored residues to construct an alanine insertion mutant. For example, the mutant DHFRcoli-1A2 contains an inserted alanine residue between the first and the second residues. When the n^th^ residue is an alanine, (n-1)An and nA(n+1) have the same sequence. Therefore, the actual number of the mutants was 145 since DHFRcoli contains 13 alanine residues. Again, we computed the 3D structural models of these 145 DHFRcoli mutants with both AlphaFold2 and RoseTTAfold. When the n^th^ residue is an alanine residue, the same computational and experimental model was used for the mutants (n-1)An and nA(n+1). For simplicity, we refer them as 158 alanine-insertion mutants in this paper.

Similar to the circular permutants, the distribution of the backbone RMSD values of the models of these 158 alanine-insertion mutants showed that AlphaFold2 generated near-native 3D models (Figure 4a & Table S2). However, the unfoldable mutants had a significantly larger mean RMSD value than the foldable mutants (Figure 4b). When we divided the models into two groups based on their RMSD values relative to the RMSD value of DHFRcoli-1A2, the mutants in the Low RMSD group had higher foldability than the mutants in the High RMSD group (Figure 4c). The mutants in the High RMSD groups formed local clusters as well (Figure 4d). Among these 158 mutants, there were 79 (50.0%) mutants with an alanine residue inserted either within secondary structural elements or out of secondary structural elements respectively. As shown in Figure 4e, among the 79 mutants with the insertion site within secondary structure units, 23 (29.1%) were foldable and 56 (70.9%) were unfoldable. The 79 mutants with the insertion site out of secondary structure units, 64 (81.0%) were foldable and 15 (19.0%) were unfoldable. Therefore, alanine insertion within secondary structure units resulted in unfoldable mutants with higher probability. When the insertion site was within secondary structure units, there were significantly more foldable permutants in the Low RMSD group than that in the High RMSD group (20 v.s. 3). Similarly, when the insertion site was out of secondary structure units, there were also more foldable permutants in the Low RMSD group than that in the High RMSD group (59 v.s. 5). When the secondary structure rule and the RMSD rule were combined, the percentage of foldable permutants in the Low RMSD group (40.0%, or 20/50) was higher than the percentage of the foldable permutants in the High RMSD group (10.3%, or 3/29). When the insertion site was out of secondary structure units, these percentages were 86.8% (59/68) and 45.4% (5/11). Therefore, no matter where the insertion site is, the mutants in the Low RMSD group has higher foldability than those in the High RMSD group. These data indicated that the conclusions from the circular permutants above are also valid for alanine insertion mutants. In addition, there was a positive correlation between the RMSD values of the circular permutants and the alanine insertion mutants (Figure 4f), confirming the experimental data showing that alanine insertion and circular permutation were different but comparable on DHFRcoli ^14^.

**Figure 4.**
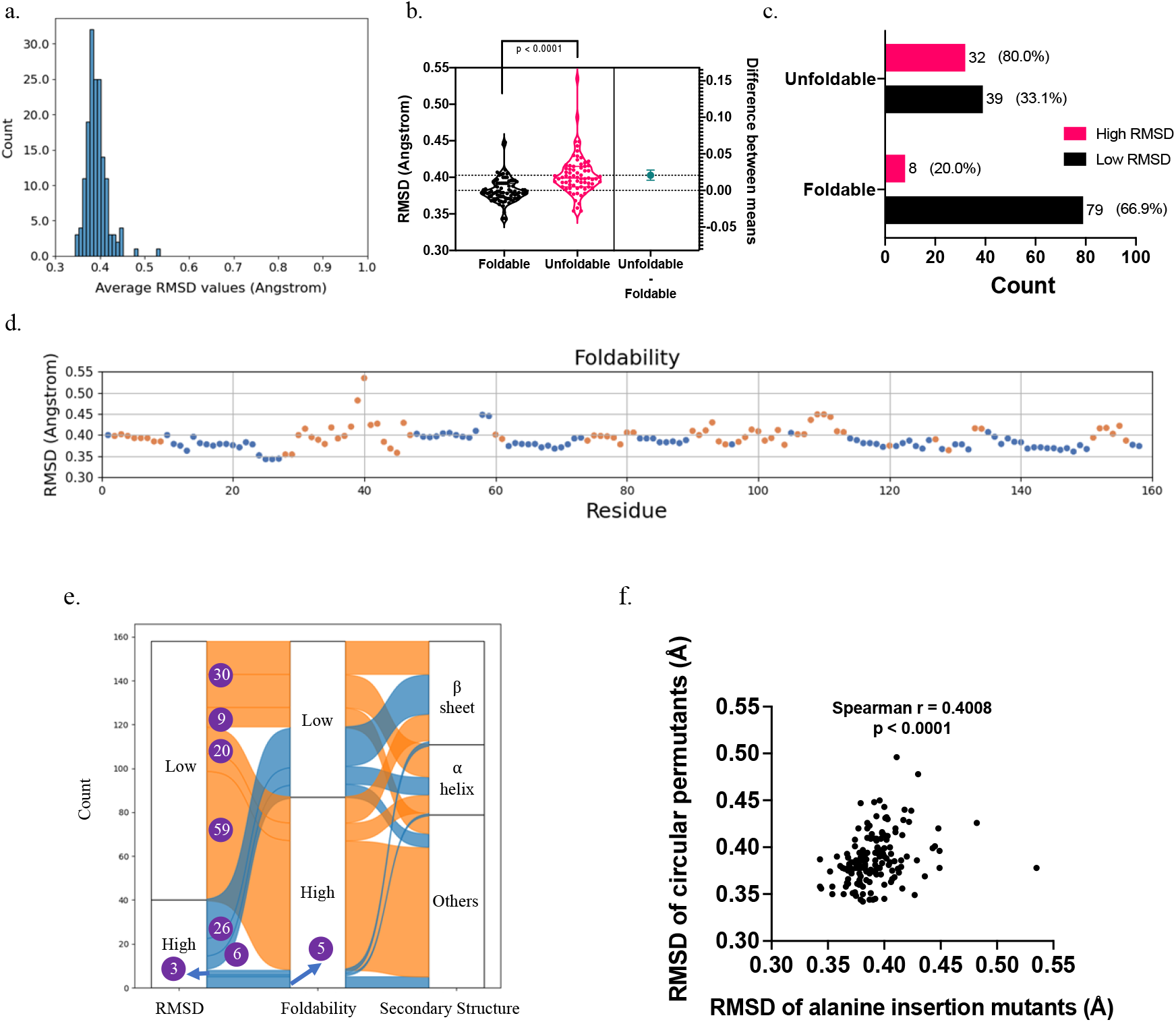
The statistics of the AlphaFold2 models of 158 alanine insertion mutants of DHFRcoli. (a) The distribution of the RMSD values of the 158 AlphaFold2 models. (b) The comparison of the AlphaFold2 models’ RMSD values of experimentally foldable and unfoldable mutants. The statistical p value was calculated with the unpaired t-test. (c) The amounts of the foldable and unfoldable mutants in the Low RMSD group and the High RMSD group. The numbers beside the bar are the mutant number and the percentage in the corresponding group. (d) The RMSD distribution of the 158 DHFRcoli alanine insertion mutants. The blue dots indicate the experimentally foldable mutants, and the orange dots indicate the unfoldable ones. (e) The correlation between RMSD values of the models, the foldability of the permutants, and the secondary structure units of the residues. The numbers in purple circles are the numbers of the permutants in different categories. (f) The correlation between the RMSD values of the AlphaFold2 models of the circular permutants and the alanine insertion mutants.

Different from the circular permutants, RoseTTAfold generated models resembling the wild-type structure for all alanine insertion mutants (Figure 5a & Table S2), likely due to the fact that the insertion of one alanine residue had very limited consequence on multiple sequence alignment. The conclusions from AlphaFold2 held true for RoseTTAfold, including: (1) The unfoldable mutants had a significantly larger mean RMSD value than the foldable mutants (Figure 5b); (2) The mutants in the Low RMSD group had higher foldability than the mutants in the High RMSD group (Figure 5c); (3) The mutants in the High RMSD groups formed local clusters (Figure 5d). As shown in Figure 5e, when the insertion site was out of secondary structure units, the percentage of foldable permutants in the Low RMSD group (88.2%, or 15/17) was higher than the percentage of the foldable permutants in the High RMSD group (79.0%, or 49/62). When the insertion site was within secondary structure units, no mutant in the Low RMSD group was foldable due to the very limited sample size (4), and the foldable mutants in the High RMSD group were only 30.7% (23/75). Despite of this tiny discrepancy, we could see that the results from the RoseTTAfold models supported the conclusions from the AlphaFold2 models. In addition, the comparison of AlphaFold2 models and RoseTTAfold models showed that their model qualities were correlated but also different. Meanwhile, the AlphaFold2 models had lower RMSD values than the RoseTTAfold models. Therefore, the correlation between conformational variations and foldability seemed to be independent of model quality as well.

**Figure 5.**
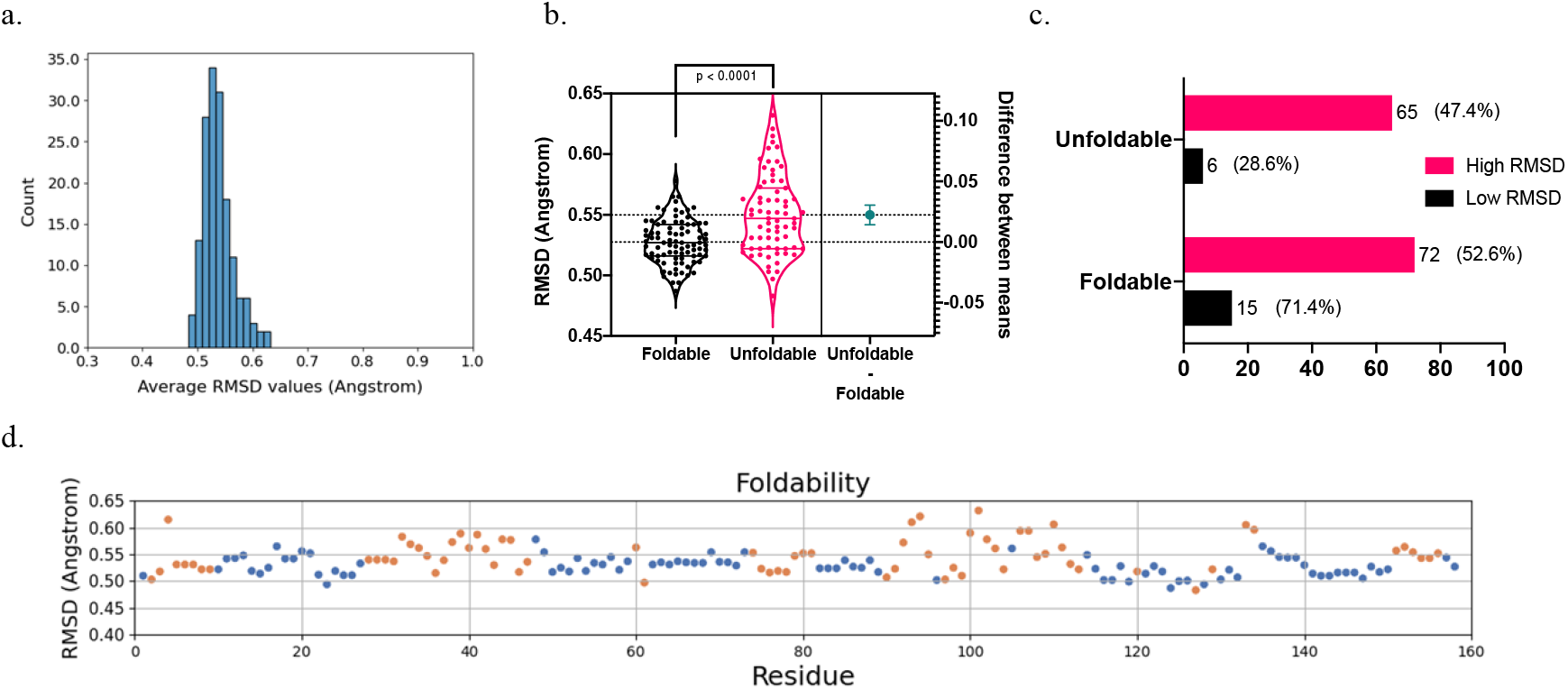
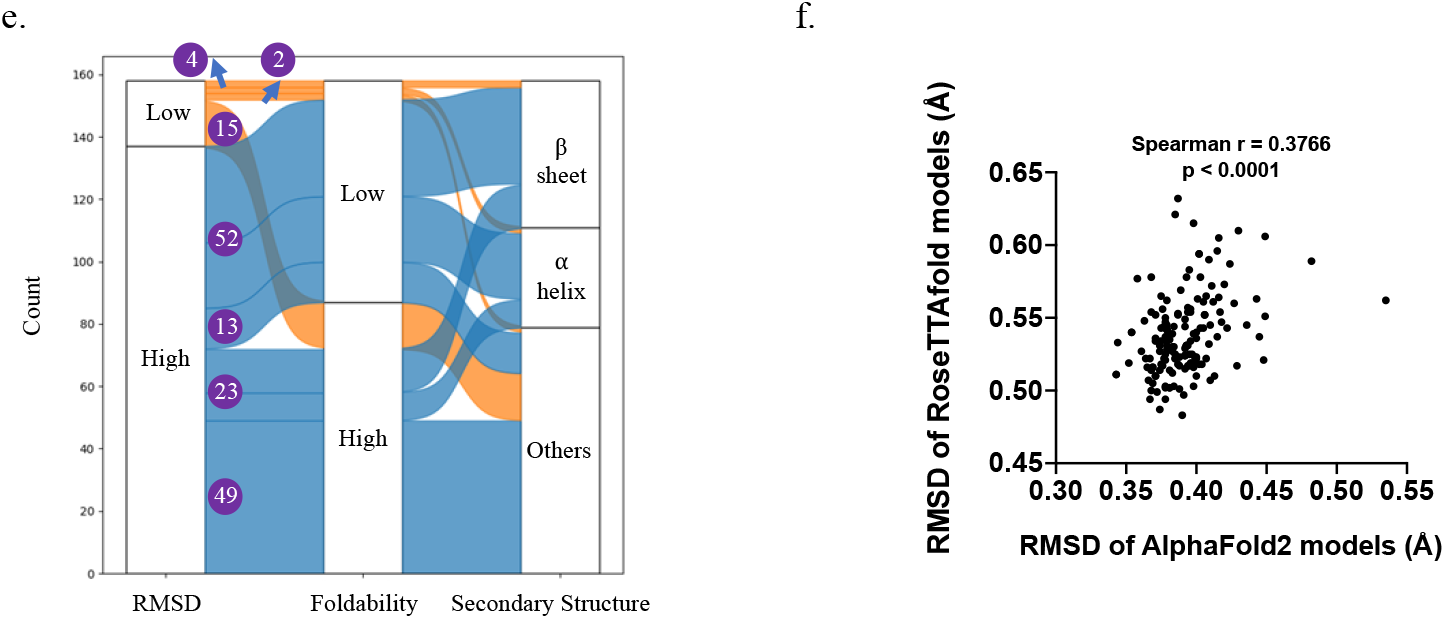
The statistics of the RoseTTAfold models of 158 alanine insertion mutants of DHFRcoli. (a) The distribution of the RMSD values of the 158 RoseTTAfold models. (b) The The comparison of the RoseTTAfold models’ RMSD values of experimentally foldable and unfoldable mutants. The statistical p value was calculated with the unpaired t-test. (c) The amounts of the foldable and unfoldable mutants in the Low RMSD group and the High RMSD group. The numbers beside the bar are the mutant number and the percentage in the corresponding group. (d) The RMSD distribution of the 158 DHFRcoli alanine insertion mutants. The blue dots indicate the experimentally foldable mutants, and the orange dots indicate the unfoldable ones. (e) The correlation between RMSD values of the models, the foldability of the permutants, and the secondary structure units of the residues. The numbers in purple circles are the numbers of the permutants in different categories. (f) The correlation between RMSD values of the AlphaFold2 models and the RoseTTAfold models of the DHFRcoli alanine insertion mutants.

### Conformational variations quantitatively correlate with protein foldability

We then asked to what extent the conformational variations of computational models quantitatively correlate with protein foldability. In the alanine insertion study, Shiba et al. ^14^ reported the precipitant ratios of all mutants from E. coli cell lysates. We found that the RMSD values of the computational models were quantitatively correlate with the precipitant ratios with a Spearman r being 0.5421 (p < 0.0001) for the AlphaFold2 models and a Spearman r being 0.3554 (p < 0.0001) for the RoseTTAfold models (Figure 6), indicating that the RMSD values could be a prediction of protein foldability. Interestingly, for foldable circular permutants, Iwakura et al. ^11^ obtained their in vitro enzymatic activity data (k_cat_ and K_M_) and conformational stability data (deltaG and m-Value), but there were no clear correlations between the RMSD values and these data (Figure S3a). On the contrary, even only foldable alanine mutants were included, a positive correlation between their RMSD values and the precipitant ratios was noticed for AlphaFold2 models (Figure S3b). These data indicated that the RMSD values of the predicted models are correlated with proteins’ native foldability but not their in vitro activities. Nonetheless, this would need further investigations since circular permutation and alanine insertion are comparable but not exactly equivalent ^14^.

**Figure 6.**
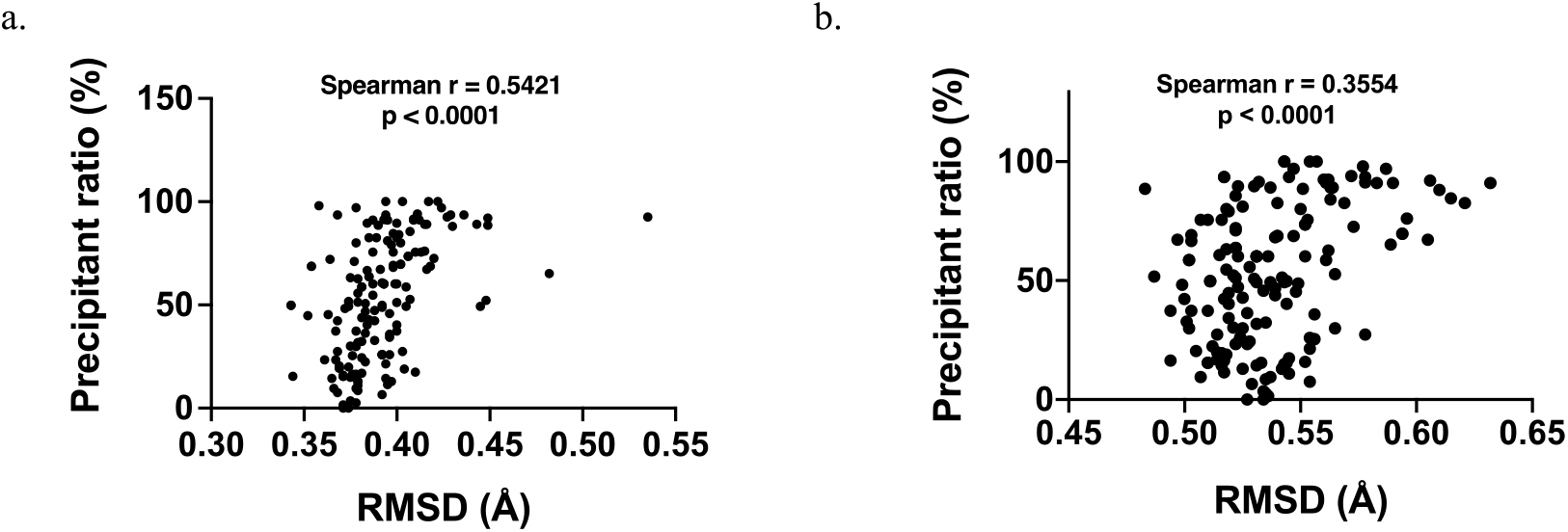
The quantitative correlation between the RMSD values of computational models and protein foldability. (a) Plot for the AlphaFold2 models of the DHFRcoli alanine insertion mutants. (b) Plot for the RoseTTAfold models of the DHFRcoli alanine insertion mutants. The precipitant ratios were obtained from ^14^.

## Discussion

The protein folding problem is a “holy grail” problem in biology. To predict the natural 3D structure from a sequence of amino acids has been a tantalizing but extremely challenging task in the last 50 years. With the assembly of the biannual Critical Assessment of Structure (CASP) meetings, global efforts have pushed forward the progress in the computational prediction of protein structure ^16^. At the recent CASP14 meeting, artificial intelligence (AI) based AlphaFold2 ^4^ and RoseTTAfold ^5^ achieved high-accuracy prediction of protein structures comparable to experimental data. However, in a recent example, only a small fraction of computationally designed proteins could be expressed as well-folded proteins ^6^.

A possible reason of this caveat of the current AI-based protein structure prediction tools is that from the training of PDB data, only the protein folding code could be extracted. That is, neither AlphaFold2 nor RoseTTAfold could directly obtain the kinetics information of protein folding. Meanwhile, the folding of a protein is not only determined by whether it could be theoretically folded, but also affected by its folding kinetics ^2^. If a peptide chain could not be folded in a suitable time scale, its folding would be trapped by frustration and fails to form folded structures. Therefore, it is not surprising that the current AI-based prediction tools could not determine the biological foldability of a protein (Figure 1 & 4).

Nonetheless, considering that the structures in the PDB database are intrinsically foldable, it is possible that the network properties of AI-based methods contain some information of protein foldability. For example, the residue-residue contact information from multiple sequence alignment might contain foldability information, since unfoldable mutants have been discarded by evolution. Then a reasonable inference is that there is some foldability information hidden in the computationally folded models. The question is how that information could be extracted.

Circular permutation modifies the termini of a protein but does not change its sequence composition. Most circular permutants could fold into native structures, although their folding kinetics might change. The alanine insertion method is more conserved on perturbing protein sequence. Since the protein sequence is minimally changed, the computational tools would be hard to tell the difference on the foldability of different constructs. Our data showed that this is true, since AlphaFold2 and RoseTTAfold predicted near-native 3D models for both foldable and unfoldable DHFRcoli permutants and alanine-insertion mutants. However, it was interesting to notice that the conformational variation of the computational models contains some information on the foldability of a protein in our work.

A caveat of our work is that the experimental data only reflected the foldability of a peptide sequence kinetically accessible within a laboratory time scale. For example, Iwakura et al. ^11^ refolded some permutants from inclusion bodies in vitro. But this should not be a big problem, since a protein not foldable within a reasonable time scale might be not biologically useful. Another question is if the conclusions from DHFRcoli in this work could be generalized to other proteins, which would need future investigations. It also remains to see if the other AI-based protein prediction methods would have similar results.

Our work would be helpful to the design of protein circular permutants and fragment complementary proteins. We suppose that the conclusions in this work could be also useful in the de novo design of soluble proteins. Based on our findings, we hypothesize that including nonfoldable protein sequences in the training data of neural networks would be useful for the AI-based prediction methods to predict protein foldability. In this regard, we believe that publishing unsuccessful protein design data is scientifically valuable ^17^. Lastly, we hope that our work would be a hint to the establishment of AI-based prediction of protein folding kinetics in the future.

## Author contributions

S.L. conceived the idea and did the computational work. K.W. and C.C. acquired the experimental data of the mutants. S.L. and K.W. analyzed and interpreted the data. S.L. wrote the manuscript. All authors reviewed and approved the submitted manuscript.

## Additional information

### Competing financial interests

The authors declare no competing financial interests.

## Acknowledgements

We would like to thank the support from the other members of our lab and Prof. Yongqi Huang for kind suggestions. S. L. was supported by the grants from National Natural Science Foundation of China (31971150), Department of Science and Technology, Hubei Provincial People’s Government (2019CFA069), and Hubei University of Technology.

## Methods

### Construction of the sequences of the circular permutants

The circular permutated sequences were constructed according to the construction methods described in ^10,11,18^. A five-glycine (GGGGG) sequence flanked the C-terminal residue (residue 159) of the wild-type sequence, connecting the downstream sequence in the permutated protein. When the N-terminal residue was not methionine (M), the extra M was added as required in protein expression. However, as describe in ^18^, an extra M was also added for the DHFRcoli-20 sequence since this was done in protein expression to protect the removal of M20 by the methionyl-aminopeptidase.

### Construction of the sequences of the alanine insertion mutants

As describe in ^14^, an alanine insertion mutant had an alanine residue inserted between two neighbored residues. For the 195-residue DHFRcoli, there would be 158 insertion sites. However, When the n^th^ residue is an alanine, (n-1)An and nA(n+1) have the same sequence. So the actual number of the alanine insertion mutants were 145 since DHFRcoli contains 13 alanine residues.

### Acquire of the experimental data

The experimental data of the circularly permutants were obtained from the reference ^11^. From the original paper, we could not determine the relative expression level (solubility) of the permutants, so we assigned foldability (foldable vs unfoldable) according to the enzymatic activity data and the CD data. The experimental data of the permutant 54 (DHFRcoli-54) was missing, but it was assigned as foldable since this position was not included in any folding element ^11^. The experimental data of the alanine insertion mutants were obtained from the reference ^14^. The alanine insertion mutant 67 (DHFRcoli-67A68) did not have the precipitant ratio data, but this mutant was assigned as foldable in this work, since it was not included in any folding elements. The data were extracted from the figures using WebPlotDigitizer (https://automeris.io/WebPlotDigitizer).

### Protein structure prediction with AlphaFold2

The 3D structures of the DHFRcoli sequences were predicted with AlphaFold2 using MMseqs2 v1.2 ^19^. Templates were used, and the Amber force field was used to relax the model structures. The multiple sequence alignment (MSA) mode was MMseqs2 (UniRef+Environment), and the pair mode was set as “unpaired+paired”. The recycle number was three. For each sequence, five unrelaxed and five relaxed models were generated, and the relaxed models were used in followed analyses.

### Protein structure prediction with RoseTTAfold

The 3D structure sof the DHFRcoli sequences were predicted with the RoseTTAfold using the end-to-end version. The checked out main version was 20210803 with UniRef30 HHsuite (2020.06), BFD (id30_c90), and pdb100 (2021Mar03). For each sequence, five models were generated and used for analyses.

### RMSD calculation

The RMSD values between the predicted models and the crystal structure (PDB ID 1RX4) were calculated with the “align” function in Pymol ^20^. Only backbone non-hydrogen atoms (CA+CB+C+O) were used in the alignment. The aligned structures were prepared in Pymol and the models were colored by pLDDT scores in spectrum.

## Notes

### Competing Interest Statement

The authors have declared no competing interest.

